# Sexually Dimorphic Visual Processing in the Yellow-Fever Mosquito, *Aedes aegypti*

**DOI:** 10.64898/2026.09.02.748965

**Authors:** Carlos Ruiz, S. David Stupski, Ruchao Qian, Amber Wu, Sophia Dominguez, Jamie C. Theobald, Jeffrey A. Riffell

## Abstract

Flying insects process motion across the visual scene to regulate their ground speed, correct course deviations, and respond to behaviorally relevant stimuli such as avoiding predators or finding hosts. The yellow-fever mosquito *Aedes aegypti* provides a useful model for studying these visually guided behaviors, as both sexes use vision to track moving features. Approaching a moving target presents mosquitoes with a demanding visual problem: they must extract target motion from the changing visual scene while simultaneously controlling their own flight and responding to potentially dangerous movements. To understand this, we first studied free-flight responses to static and moving vertical targets in a wind tunnel system capable of projecting dynamic visual stimuli, and found that males and females are sexually dimorphic with respect to their responses to target motion. When the feature was static, both males and females advanced toward it during flight; however, when it oscillated laterally at 2 Hz, only females continued to pursue it. In a subsequent yaw-unrestrained magnetic tethering paradigm, we showed that this divergence cannot be simply explained by sex-specific differences in visuomotor tuning dynamics, while micro-CT reconstructions revealed only modest sexual differences in eye morphology and broadly similar patterns of estimated spatial acuity. Together, these results indicate that males retain the biomechanical capacity to track oscillating targets but actively suppress approach behavior during free flight, whereas females pursue salient features independently of target motion. This pattern aligns with sex-divergent visual ecologies because the risk of approaching a large moving feature too closely likely outweighs the benefits for males seeking mates, whereas for host-seeking females the reproductive necessity of blood feeding overrides this avoidance.

## INTRODUCTION

Insects rely on vision to perform complex navigational tasks and complete their life cycles. For a flying insect, this may include associating odor sources with visually salient features in the environment (Blake and Riffell, 2025; Van Breugel et al., 2015; Stupski and Van Breugel, 2024), avoiding collisions in high-density aggregations (Singh et al., 2024), and maintaining biomechanical stability to compensate for unexpected optic flow (Fuller et al., 2014; Ruiz and Theobald, 2021, 2020). How insects process motion along their retinas and which visual features elicit determined responses, are fundamentally linked to their ecological niches. For example, subspecies of flies isolated in different deserts navigate toward features whose height matches that of the host cacti in their respective habitats (Rimniceanu et al., 2024). Within a single species, divergent ecologies and life histories between sexes can also impose sexually dimorphic evolutionary pressure on sensory systems. Male house flies, for instance, have a specialized array of photoreceptors tuned to detect the motion of females during mating pursuits, commonly referred to as the “love spot” (Burton et al., 2001; Burton and Laughlin, 2003).

Mosquitoes as a group are another example where each sex has considerably different visual life histories. Although male and female mosquitoes share many navigational goals, such as avoiding looming threats (Cribellier et al., 2024) or finding nectar sources to replenish carbohydrate stores (Dexter, 1913; Lahondère et al., 2020), each sex also has highly sex-specific navigational tasks that require specialized sensory processing. For male mosquitoes, visual processing is modulated by acoustic cues in order to intercept a mate, while avoiding other individuals in an aggregation (Gupta et al., 2024); there is no evidence of a similar phenomenon in females. For females, vision also guides highly sex specific behaviors, such as finding an oviposition site (Bernáth et al., 2008). However, in terms of life history a female mosquito’s necessary pursuit of a host is an especially sexually dimorphic visually guided task.

Host landing sites, however are rarely stationary environmental features (Matherne et al., 2018) and may carry fundamentally different risks and rewards for each sex. For female *Ae. aegypti*, a moving feature may be representative of a viable blood meal, an essential requirement for completing the gonotrophic cycle (Beklemishev, 1940; Valzania et al., 2019). While there is suggestive evidence that moving hosts are more attractive to female *Ae. aegypti* than stationary ones (Sippell and Brown, 1953), motion is usually accompanied by other cues that females may find aversive, such as air currents generated by the host’s defensive movements (Matherne et al., 2018). Males, on the other hand, typically navigate toward tall static features when foraging or seeking perches. In contrast with females, males do not tend to approach hosts directly but gather near them in loose aggregations where they find and intercept blood-seeking females. While this behavior is considered a strong component of mating success (Hartberg, 1971; Jones and Pilitt, 1973), approaching a large, active host too closely exposes males to potentially lethal risks (Vinauger et al., 2018; Cribellier et al., 2024). Because moving visual targets carry different risk-reward significance for each sex, here we investigate whether the motion of an ersatz host carries differential behavioral valence between mosquito sexes.

We begin by quantifying free-flight navigation of male and females mosquitoes towards static and moving vertical targets projected in a wind tunnel system capable of tracking mosquitoes’ position and heading. In this system, retinal slip during approach to a stationary target results exclusively from self-motion, whereas the motion of an oscillating target is the combination of the mosquitoes’ translation in space and the motion of the target itself. In dipterans, visual fixation strategies rely heavily on gyroscopic feedback from halteres (Rimniceanu et al., 2023; Sherman and Dickinson, 2004; Fox et al., 2010), the club-shaped sensory organs that evolved from dipteran hind wings (Weatherbee et al., 1998), we complemented free-flight assays with yaw-unrestrained magnetic tethering to isolate any potential differences in visuomotor tuning between sexes, while preserving the haltere derived component that influences bar fixation. This approach allows us to determine whether sexually dimorphic steering responses result from active behavioral modulation rather than underlying constraints, either biomechanical or in visuomotor processing. While much work has been done to unravel the evolution of navigation strategies in closely-related species under different ecological pressures (Rimniceanu et al., 2024; Takagi et al., 2024; Prieto-Godino et al., 2017), less is known about how each sex’s brain accomplishes dimorphic computations for varying navigational goals within a species (Nordström et al., 2008; Berg et al., 2025).

## MATERIALS AND METHODS

### Experimental Animals

*Ae. aegypti*: Liverpool strain mosquitoes (LVP, BEI Resources) were maintained in a laboratory colony at the University of Washington. Experimental mosquitoes were reared in mixed-sex chambers at 28° C under a 14 h light, 10 h dark cycle. Mosquitoes had *ad libitum* access to 10% sucrose solution prior to experiments. All experiments were performed on co-housed individuals aged 4 to 8 days and took place during the dusk phase of their photoperiod.

### Behavioral Wind Tunnel for Free flight Experiments

To characterize visually guided navigation in free-flying mosquitoes, we built an opaque white acrylic wind tunnel (0.3 m x 0.3 m x 0.9 m). A transparent top panel on the test section allowed visual stimuli projection and behavioral recording using an externally mounted projector and cameras. We measure the air speed at the center of the test section with an OMEGA HHF1000 Series anemometer and maintained it at 0.15 m/s (sd=0.0026 m/s, turbulence intensity=1.71%) across trials. Mosquitoes were released into the system by attaching a rearing container to the underside of the containment section of the system and opening a trap door until 5-10 individuals entered. While confined in the antechamber, individuals were exposed to 5 s pulses of 2% CO_2_, every 5 minutes to prime flight behavior (Alonso San Alberto et al., 2022). CO_2_ was released upwind from behind the projection screen through evenly spaced nozzles to ensure homogeneous distribution across the test section rather than a structured plume, with concentration and pulse volume regulated by two mass flow controllers (ALICAT MC-200SCCM-D). Primed mosquitoes were given 30 minutes to enter the test section of the arena by flying upwind through a 6 cm diameter hole in a dividing PTFE mesh, remaining individuals were vacuumed out before the next batch was released.

### Video Recording and Image Processing in Wind Tunnel Assays

We recorded mosquito flight in the test section at 140 Hz, 1.5 ms exposure time, 1440x1080 px resolution, using two synchronized Flir Blackfly S USB3 cameras (BFS-U3-16S2M-CS) fitted with IR-pass filters (*>*850 nm) mounted above the arena in a stereoscopic array. The test section was back-illuminated with diffused infrared arrays (*∼* 850 nm) located beneath the floor (*∼* 90 W), and behind side panels (*∼* 40 W), to maximize silhouette contrast. Recording and stimulus onset were automatically triggered by motion detection. We trained two DeepLabCut (DLC) (Mathis et al., 2018) models to independently annotate head and abdomen positions in male and female videos. To ensure accuracy, we visually verified DLC marker placement against raw footage for each trajectory, and defined usable length as the number of consecutive frames spanning from trial onset until a disruptive event (typically wall collisions resulting in erratic flight). Curated 2D annotations were processed through a custom OpenCV (Bradski, 2000) pipeline to generate 3D reconstructions.

### Visual Stimuli in Free-flight

Visual stimuli were projected onto a white PTFE mesh located on the anterior wall of the test section using a halogen lamp projector (BENQ TK700-STi) operating at 60 Hz refresh rate. The beam was physically masked to illuminate only the projection screen, making it the only source of visible light in the system. The visual target was a tall black bar (60x160 mm) on a white background, projected on the anterior wall of the arena and subtending 12° horizontally and 31° vertically, when viewed from the tunnel entrance. The bar was randomly located at the start of each trial 100 mm to the left or right of the vertical midline of the screen and its position logged. No-bar trials contained a stationary ‘ghost’ bar that was white and provided no contrast against the white background. During moving trials the bar center moved sinusoidally at 2 Hz by approximately ±33 mm. Motion was set to start 0.36 s (50 frames) after mosquito detection and lasted for 6 cycles.

### Analysis of Free-flight Data

We defined mosquito position as the midpoint between head and abdomen markers, with heading as the unit vector from abdomen to head. Trajectories were smoothed using a fourth-order Butterworth filter with a 15 Hz cutoff to minimize reconstruction jitter before derivative analyses. Statistical comparisons were restricted to tracks spanning at least 75% as long as the window of interest, unless otherwise stated. We quantified approach toward the visible or ghost bar by calculating the mean distance between the mosquito’s position and bar center during a late time window in the trial (about 1.43-2.14 s), which captures stimulus-driven responses, while avoiding near-screen maneuvers. We analyzed this metric using a two-factor ANOVA evaluating sex, treatment, and interaction, followed by seven planned two-sided permutation contrasts (10 000 permutations) and report Holm-corrected p-values for multiple comparisons. To evaluate whether mosquitoes perform compensatory steering in response to bar motion, we analyzed the amplitude spectrum of the lateral velocity for all trajectories across treatments during motion presentation (about 0.36-3.36 s). To isolate the effect of the motion of the target on mosquitoes’ flight trajectories, we compared the excess spectral amplitude in the 1.6-2.4 Hz band, in the frequency domain of the mosquito’s lateral velocity across sexes and treatments. We fitted permutation-based linear models with sex, treatment, interaction, and log(track length) as predictors. Planned-contrast *P* values were corrected using the Benjamini-Hochberg false discovery rate procedure.

For frontal fixation analysis, we define error angles as the angular error between the mosquito’s heading and a vector from head to bar center in each frame. Statistical comparisons of body orientation were run on full trajectories after excluding the first 0.36 s during which the bar remained stationary in all treatments. Analyses retained only frames in which mosquitoes were within the upwind two-thirds of the arena (*≤* 200 mm from the entrance). Differences across treatments were evaluated using the fraction of frames in which the bar center fell within *±* 15^*°*^ of the mosquito’s forward body axis, and analyzed using a two-factor ANOVA with sex, treatment, and their interaction as fixed effects, followed by nine pairwise comparisons using two-sided Welch’s t-tests. P-values presented were adjusted using the Benjamini-Hochberg correction.

### µCT Scanning and Image Reconstruction

To compare the general morphology across sexes and evaluate potential differences in facet diameter and spatial acuity, we scanned heads of male and female *Ae. aegypti* mosquitoes using micro-computed tomography (µCT) to examine eye characteristics in three dimensions. We first anesthetized the animals by cold exposure, and heads were removed and stored in 70% ethanol for at least 24 h before staining. To enhance the contrast of soft tissues under X-ray, we stained all samples with 5% phosphotungstic acid (PTA) solution dissolved in 70% ethanol. To prevent tissue deformation during scanning, all samples were chemically dehydrated before imaging by transferring the stained heads through a graded series of 80%, 90%, and 100% ethanol. After dehydration, we immersed the samples in hexamethyldisilazane (HMDS) for 24 h to replace the ethanol and stabilize internal structures. We then left the samples in sealed microcentrifuge tubes to allow gradual evaporation of HMDS before scanning. This drying procedure reduced internal movement in samples and minimized deformation artifacts during µCT imaging. Scans were acquired at the University of Florida Nanoscale Research Facility using a ZEISS Xradia 620 Versa 3D X-ray microscope. The volumetric datasets were automatically reconstructed using the built-in reconstruction pipeline of the scanner. Although we conducted quantitative Ommatidia Detecting Algorithm (ODA) analysis (Currea et al., 2023) on one specimen per sex, we visually examined all six µCT-scanned specimens and did not observe obvious differences in eye morphology among individuals. The remaining scans were retained for qualitative validation to confirm that the selected specimens were morphologically representative and the final measurements were consistent with the broader sample. We imported all reconstructed volumes into 3D Slicer (version 5.6.2) (Fedorov et al., 2012) for manual alignment to a standardized anatomical frame of reference. We removed non-relevant tissues and isolated visual structures from the rest of the head using the Segment Editor module. We exported the segmented eye data for ODA analysis (Currea et al., 2023), an automated pipeline designed to detect individual ommatidia from volumetric or image-stack data and to extract facet-level morphological information across the eye. We used ODA to calculate morphological measurements of the male and female eyes, including ommatidial positions, facet-related metrics, and spatial resolution. Automated detections were manually verified against original reconstructions to correct imaging artifacts, incomplete boundaries, or local segmentation errors. Original reconstructions were also used to correct misidentified structures or inconsistent measurements before final analysis, ensuring all final outputs were validated by manual inspection.

### Magnetic Tethering in Virtual Reality

In order to investigate whether males and females had fundamentally different visuomotor tuning profiles we designed and built a yaw-unrestrained magnetic tethering system originally developed for flies(Bender and Dickinson, 2006) and later for mosquitoes (Liu and Vosshall, 2019). For these experiments, an individual mosquito was temporarily chilled on a Peltier stage set to 6° C. The mosquito’s dorsal thorax was then fixed to the dull end of a minutien pin (Fine Science Tools, Forster City, CA) with UV-activated glue. The fine point of the mosquito-pin complex was then placed in the pivot of a low-friction V1.50 jewel bearing (Swiss Jewel Company, Bala Cynwyd, PA) attached to a bar magnet embedded in a custom 3D-printed positioning rod. The positioning rod was attached to a 3-axis micro-manipulation system to precisely place the mosquito over a stack of ring magnets mounted on a custom 3D-printed stage made from PLA. The stage allowed for a light path for video recording from underneath the floating mosquito using a computer vision camera (Basler aca640, Basler AG, Ahrensburg, Germany) with a 10X (13 - 130 mm FL) C-Mount, Close Focus Zoom Lens (Edmund Optics, Barrington, NJ) collecting images at 50 Hz (Fig 1A, B). A sampling frequency well beyond the Nyquist cutoff of any stimulus tested in this study. Illumination was handled by 950 nm infrared lighting rings mounted above the arena. All experiments were performed in the dark except for the LEDs that form the basis of the virtual reality system.

**Figure 1.**
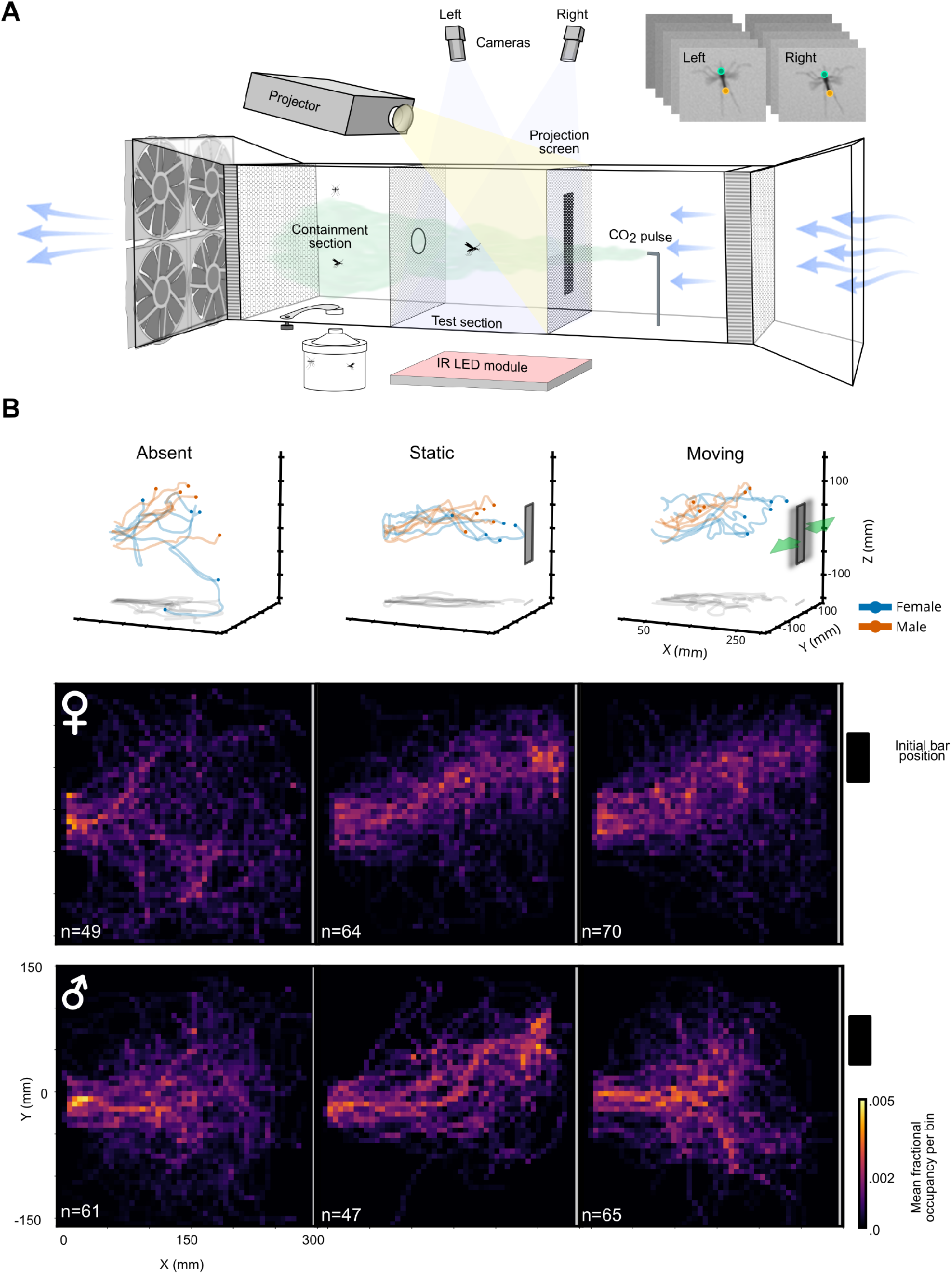
Experimental setup for studying mosquito visual responses to moving features. **A:** General overview of the free flight arena. Mosquitoes were released into a containment section at the back of the system, where individual mosquitoes are lured into a test section with CO_2_ pulses. Flight behavior is recorded using two synchronized cameras, to generate 3-D reconstructions of the trajectories and mosquito heading. **B:** Occupancy heatmaps showing the effect of the presence and 2 Hz oscillation of a visual target on the X,Y distribution of female and male mosquitoes during trials. Sample tracks show complete reconstructions of a subset of tracks per treatment for illustrative purposes. Static and moving trials contain an off-center black bar (right-side trials have been mirrored) spanning 12° from the entrance to the test section. The bar is projected on the anterior wall of the system and its initial position is represented as a black rectangle. Both males and females show similar foraging patterns of occupancy in the absence of a visual stimulus, and seem equally attracted to a static bar. However, adding a 2 Hz lateral oscillation to the bar stops males from approaching, while not having a clear effect on females.

**Figure 2.**
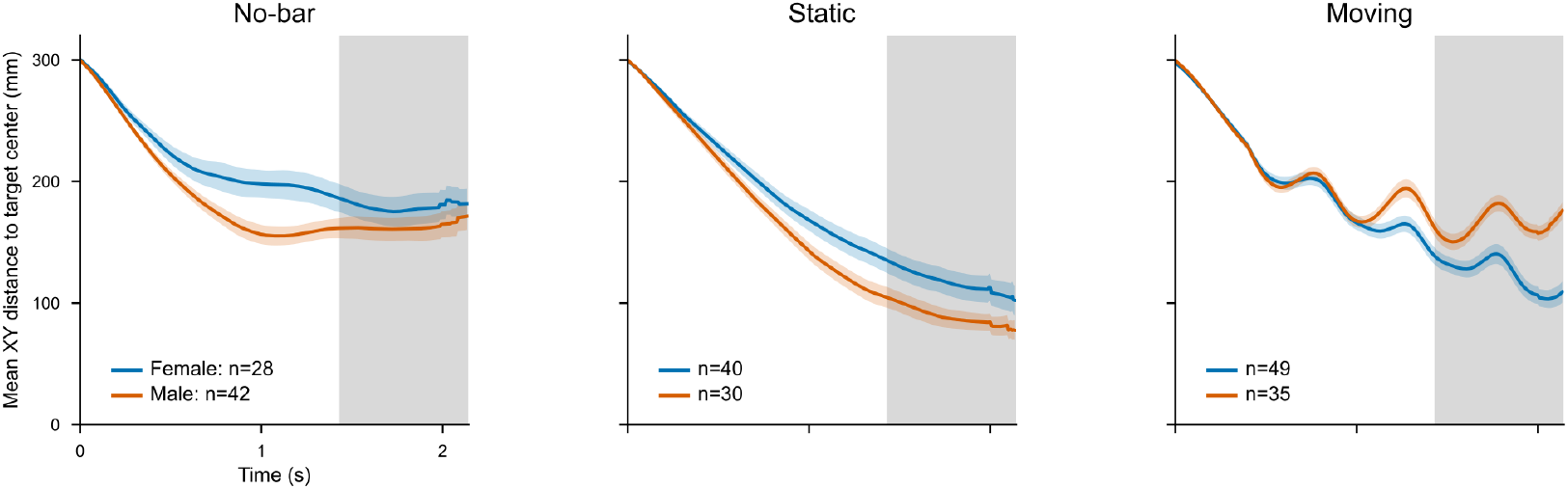
Bar motion influences male approach. Mean distance from the midpoint of the mosquito body to the horizontal center of the visible or ghost bar, for female (blue) and male (orange) *Ae. aegypti* under no-bar, static-bar, and moving-bar conditions. Solid lines and shaded envelopes represent the mean and SEM across analyzed tracks respectively. Distance to bar center was evaluated late in the trial, about 1.4-2.1 s from the start, which provided enough time for mosquitoes to be exposed and respond to the visual stimulus presented. Both sexes show substantial and similar approach to a static bar compared to the no-bar condition. However, in the presence of a moving bar, males stop their upwind advancement, resulting in a larger mean distance to the bar during the analysis window.

### Virtual Reality System

Visual stimuli were generated using the generation 3 version of the Reiser Lab modular display system (Reiser and Dickinson, 2008). Our system surrounded the mosquito preparation in a full 360° span with an array of 96 x 32 individually controllable yellow-green LEDs allowing for a 3.75° resolution along the x-axis, and is below the interommatidial angle of both male and female *Ae. aegypti* of (Muir et al., 1992). The display pattern was controlled and synchronized with image capture using a combination of Python and ROS2 Kinetic. Mosquito orientation was then processed for each individual time-stamped video frame using DLC (Mathis et al., 2018). All experiments used a star-field background that was generated by a random Poisson process where the rate parameter *λ*, determined the probability that each pixel was on and was selected as 0.125. A dark bar spanning 22.5° was projected onto a random starting position on the same star field background with only the bar moving and the star field pattern static. Wide field experiments had no bar but moved the entire panoramic starfield background.

### Experimental Design for Yaw-unrestrained Virtual Reality

Once a mosquito was inside the magnetic tethering virtual reality platform we gently exposed the animal to the experimenters breath. We breath-activated mosquitoes for two reasons: firstly, exposure to host cues increases visual tracking gains in rigid tethering preparations(Vinauger et al., 2019; Van Breugel et al., 2015) and secondly, the nature of the animal preparation necessarily brought the mosquito close to an experimenter just before behavioral experiments began, e.g. when the mosquito on the pin was inserted inside the jewel bearing facet it was brought close to the experimenter’s face. CO_2_ activation before trials was therefore used to promote flight and normalize host exposure prior to any experiment. Once the preparation was inside the virtual reality system, we then displayed motion sequences of the bar or panoramic motion and measured compensatory steering responses. The motion sequences we used were slow-to-fast chirps that spanned 60° of the 360° virtual reality arena. The chirp sequence itself lasted for 15 s and was initiated at 0.1 Hz with a final frequency of 5.4 Hz. In early experiments we tested other stimulus regimes that could be used to determine differences in visuomotor tuning differences, like sum of sines, however we found *Ae. aegypti* mosquitoes had inconsistent responses to the erratic motion relative the the fairly consistent responses to chirp sequences.

## RESULTS

### Free-flight Navigation Experiments

To investigate how each sex responds to moving features, we developed a wind tunnel system (Fig. 1A) which allowed for tracking individual mosquitoes’ flight paths from an antechamber entrance, upwind to either no projected target, a static target, or a sinusoidally moving target (Fig, 1B). Experiments without a target provide a null distribution of how mosquitoes behave in the system without the input of a salient visual feature. For experiments with a moving target, we focused on 2 Hz sinusoidal motion. 2 Hz was chosen because it was sufficiently outside of the dominant frequency of a trajectory’s lateral velocity spectrum in the absence of any visual bar, which settled around 1 Hz. This frequency was also high enough to allow multiple oscillations over the course of a mosquito’s trajectory, the mean duration of which was about 2.5 seconds.

Across all experiments, mosquitoes flew upwind, consistent with a general trend for insects to preferentially orient into the wind (Combes et al., 2023; Stupski and Van Breugel, 2024; Kennedy, 1940; Daykin, 1967). However, occupancy maps suggest a qualitative but clear distinction in the spatial distribution of mosquitoes across sexes and treatments. In the absence of a visual target, both sexes display a broad dispersion within the arena (Fig. 1B, left). When a static bar is projected, occupancy is mostly limited to a corridor connecting the entry region and the location of the visual target (Fig. 1B, middle). However, under the moving-bar condition males show a broad distribution, similar to that in the no-bar condition, whereas females retain their corridor-like distribution (Fig. 1B, right).

Mean distance to the center of the bar differed strongly among treatments and was sex-dependent. We found that, while treatment (*F*_2,220_ = 23.77, *p* = 4.52 *×* 10^*−*10^) and sex × treatment interaction (*F*_2,218_ = 9.64, *p* = 9.72 *×* 10^*−*5^) were significant, the overall main effect of sex was not (*F*_1,220_ = 0.026, *p* = 0.872). During the analysis window, females approached the visible bar, whether static (117.68 *±* 10.02 mm (mean *±* SEM)) or moving (124.37 *±* 6.93 mm), significantly more than the ghost bar in the negative control (178.13 ± 10.99 mm; static vs no-bar *p*_Holm_ = 0.0018; moving vs no-bar *p*_Holm_ = 0.0007). Males also approached a static bar (89.11 *±* 6.89 mm) significantly closer than the ghost bar (162.05 *±* 9.02 mm; difference = *−* 72.94 mm, *p*_Holm_ = 0.0007). However, in contrast to females who approached a static and a moving bar similarly (difference = 6.69 mm, *p*_Holm_ = 1.0), males got significantly closer to a bar that was static compared to one in motion (*p*_Holm_ = 0.0007). Their distance to a moving bar (164.73 *±* 5.80 mm) was no different from the no-bar condition (difference = 2.68 mm, *p*_Holm_ = 1.0). Our results indicate that target motion has a strong sexually-dimorphic impact on object approach.

### Females Dynamically Track Bar Motion in Free-flight, but Males Do Not

Crosswind mosquito flight velocity varied strongly across visual stimuli treatments (partial F = 5.47, p = 0.0004; 354 tracks), depended on sex (partial F = 2.92, p = 0.0329), and marginally on sex by treatment interaction (partial F = 3.15, p = 0.0490). To isolate whether the variation in sex and treatment reflected tracking of the moving target specifically, we used planned contrasts comparing the excess spectral amplitude of the crosswind velocity at the stimulus frequency of 2 Hz across conditions. Females showed a robust increase in the amplitude of their cross-wind velocity spectrum at 2 Hz relative to both a static bar (estimate = 15.19 mm/s, permutation p = 0.0004, BH-adjusted q = 0.0022) and a no-bar control (estimate = 18.57 mm/s, permutation p = 0.0001, BH-adjusted q = 0.0011). Female responses to a static bar were no different from the control condition, with no excess amplitude at 2 Hz (estimate = 3.38 mm/s, p = 0.4874, q = 0.6701), indicating that female steering is linked specifically to the motion of the bar, rather than an artifact of a casting frequency associated to their exposure to CO_2_ pulses (Dekker and Cardé, 2011). In contrast, male trajectories in response to a moving bar did not show the significant enrichment around the 2 Hz frequency band observed in females (Fig. 3A) and their spectrum was no different from that seen in the control or static bar conditions ((estimate vs control = 6.90 mm/s, p = 0.1352, q = 0.2124; estimate vs static = -0.62 mm/s, p = 0.9008, q = 0.9008)). These results indicate that the target’s motion did not contribute to steering responses to the same degree for males as it did for females.

**Figure 3.**
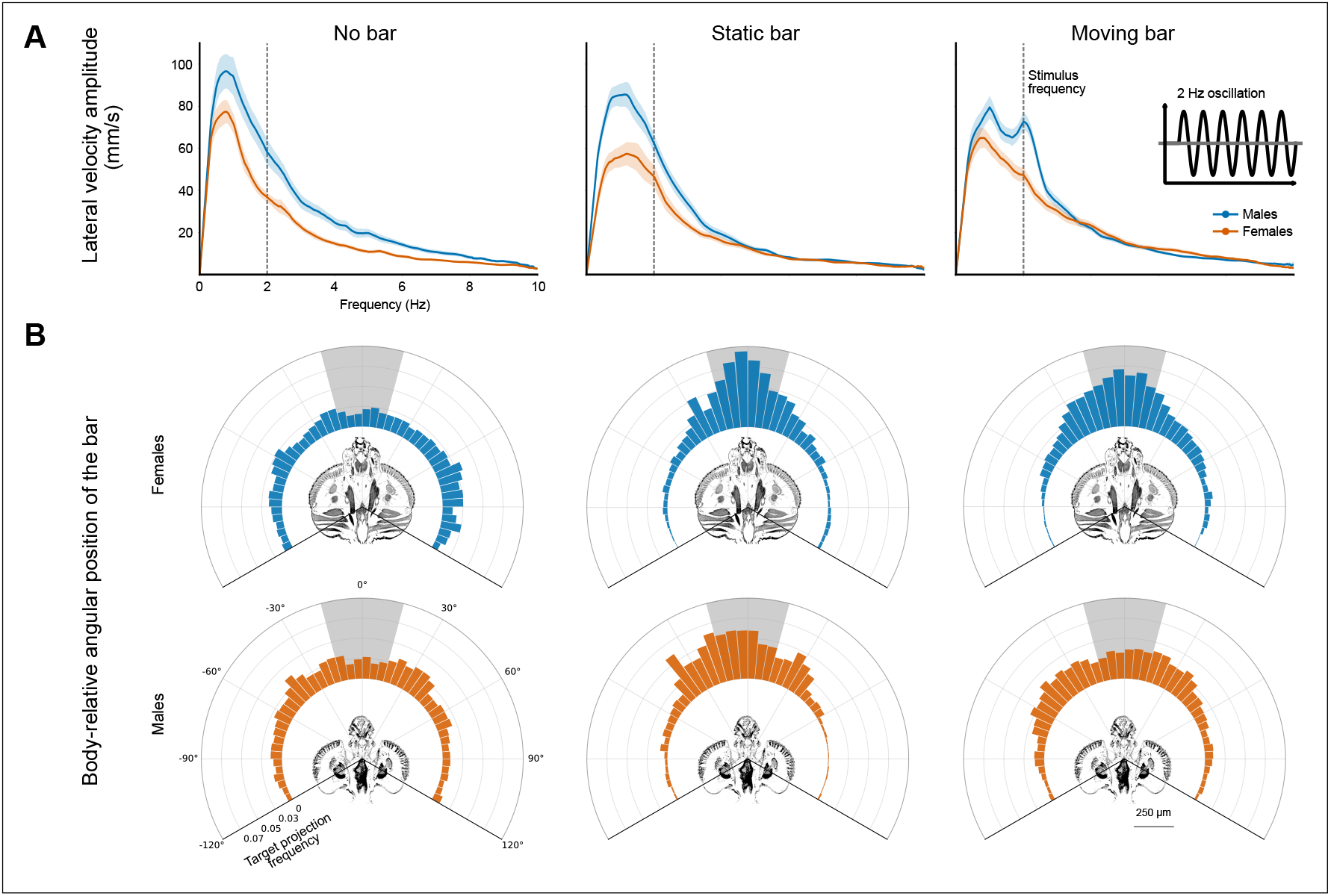
Females compensate for bar motion with a retinal-centering strategy. **A:** Sex-specific modulation of lateral-velocity spectra in response to target motion in *Ae. aegypti*. Mean amplitude of the lateral velocity frequency spectrum in male and female trajectories for the no-bar, static, and moving bar conditions. **B:** Body orientation relative to the center of the bar. Both sexes preferentially aligned with visible bars, but frontal fixation was less frequent in males when the bar oscillated at 2 Hz. Rosette histograms show the mean fraction of time in which the center of the visible or ghost bar occupies 5° bins relative to the mosquito heading. The shaded region covers *±* 15^*°*^ from the forward body axis and indicates the range used to calculate frontal fixation for statistical analysis. Rosette height represents the frequency of angular occupancy; the µCT cross-sections of the female and male heads are provided for anatomical reference.

### Male *Ae. aegypti* Use a Centering Strategy for Static, but Not Moving Bars. Females Maintain Centered Positions Regardless of an Object’s Motion

Our analysis of angular error relative to the bar using orientation data shows that frontal fixation differs significantly between sexes (two-factor ANOVA: *F*_1,350_ = 14.93, *p* = 1.33 *×* 10^*−*4^, 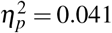) and among treatments (*F*_2,350_ = 42.79, *p* = 2.37 *×* 10^*−*17^, 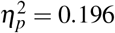), with a significant sex *×* treatment interaction (*F*_2,350_ = 6.95, *p* = 0.0011, 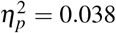). Our comparisons show that, relative to the no-bar condition (0.116), females spent on average a significantly larger proportion of time with the bar center aligned within *±* 15^*°*^ of their forward body axis during static (0.435; Welch’s *t*_92.1_ = 8.26, *q* = 6.73 *×* 10^*−*12^) and moving (0.37; *t*_112.0_ = 7.96, *q* = 6.73 *×* 10^*−*12^) bar treatments. Interestingly, frontal fixation did not differ significantly between the static and moving-bar treatments in females (*t*_121.1_ = *−* 1.51, *q* = 0.150). Similar to females, when compared to the no-bar condition (0.141), males showed robust alignment with a static bar (0.323; *t*_67.1_ = 4.42, *q* = 8.21 *×* 10^*−*5^), and a moving bar (0.204; *t*_123.7_ = 2.52), *q* = 0.0197). However, for males frontal fixation was significantly less consistent with the moving than with the static bar (*t*_66.5_ = *−* 2.90, *q* = 0.0091). While both sexes exhibited similar frontal fixation in the absence of a visual stimulus (*q* = 0.305), females showed greater frontal fixation than males with both the static (mean difference = 0.112; *t*_103.5_ = 2.22, *q* = 0.0371) and the moving bar (mean difference = 0.165; *t*_118.0_ = 5.19, *q* = 2.62 *×* 10^*−*6^).

### Differences in Steering Responses Are Not Likely Due to Visuomotor Tuning Limitations

We next asked if males’ inattention to moving features could be explained by simple differences in visuomotor tuning. In order to test whether the observed free flight behavior may be due to the inability of males to compensate for the bars motion rather than a difference in how each sex responds to motion itself, we used a yaw-unrestrained magnetic tethering system (Fig. 4A, B) designed to maximally stimulate steering responses. While not a one-to-one comparison with free-flight, we expected that if males were less capable of steering in response to moving features, unable to perceptually resolve the motion of a bar, or have differentially optimized optomotor response tuning, then they would be less proficient than females in this assay.

**Figure 4.**
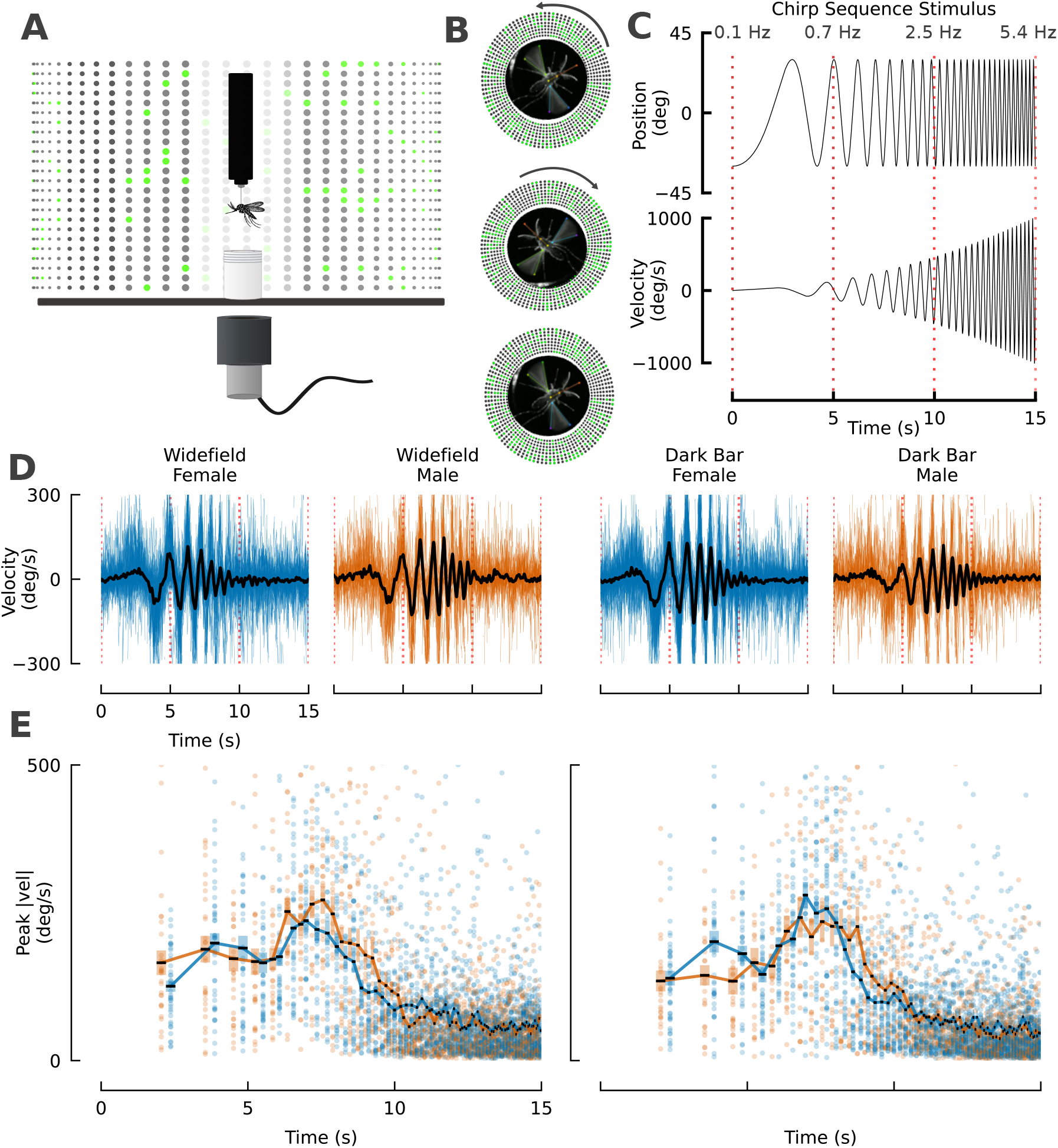
Male and Female *Ae. aegypti* mosquitoes have similar visuomotor tuning. **A:** Magnetic tethering system in which a male or female *Aedes* mosquito is glued to a minutien pin and the pointed end placed in the insert of a jewel bearing. The mosquito is filmed from below as a 360°VR system plays an input pattern. **B:** *Ae. aegypti* female performing corrective steering responses toward a 22.5 degree moving bar defined by a span of off LEDs on a star field background **C:** Position and velocity profile of a quadratic chirp sequence presented to mosquitoes. **D:** Velocity profile in response to either wide field motion (left) or a moving dark bar (right) for female (blue) and male (orange) mosquitoes. **E:** Peak velocities associated with velocity peaks of chirp sequence for males and females from both wide field and dark bar motion. n = 42 traces N = 16 Females, n = 30 traces, N = 13 males

Our input stimulus used an accelerating chirp sequence to characterize male and female steering responses to both panoramic motion, designed to maximally stimulate steering, and to moving dark bars (Fig. 4C). Using this system, we quantified the visuomotor tuning range of male and female *Ae. aegypti*. Prior rigidly tethered work shows mosquitoes will respond to moving features in similar virtual reality setups (Vinauger et al., 2019; Gupta et al., 2024). The motion was defined by a quadratic chirp sequence that accelerated from 0.1 Hz to 5.4 Hz over a period of 15 seconds (Fig. 4C).

In contrast to free-flight experiments, both sexes displayed corrective steering responses to panoramic motion and the motion of a dark bar stimulus. In virtual reality, both sexes exhibited similar dynamic tuning ranges, with no statistical difference in where the maximum steering responses occurred for either males or females (Widefield peak 1.61 Hz females, 1.71 Hz males, p=0.51, Mann-Whitney U). Maximum peak turning response velocities were also not statistically different between males and females for both wide field (females 236.7 deg/s *±* 16.3, male 271.8 *±* 24.0, p = 0.113, Mann-Whitney U) and a dark bar (female 279.9 deg/s *±* 22.6 males 241.9 deg/s *±* 34.5 deg, p = 0.524, Mann-Whitney U). Males tended to have elevated responses relative to females in the 1.7 -2.5 Hz range (Female 140.3 deg/s *±* 11.9, males 185.7 deg/s *±* 16.2, p = 0.0296, Mann-Whitney U), but were similar to females in the range for moving dark bars (females 147.6 deg/s *±* 9.9 males 181.9 deg/s *±* 21.5 p = 0.260, Mann-Whitney U). While there were marginal differences in the visuomotor dynamics, males and females showed similar biomechanical compensation over the tested tuning range, suggesting that differences in free-flight are not due to a parsimonious explanation that males are simply less able to biomechanically compensate and steer toward a moving bar or large differences in visuomotor tuning between males and females.

## DISCUSSION

In our system, mosquitoes primed with CO_2_ consistently orient themselves toward a dark static vertical bar and approach it, regardless of their sex. However, males arrest their upwind advancements when the bar is moving, highlighting a fundamentally different navigational strategy in each sex. We additionally show that this difference in behavior is not likely due to visuomotor tuning differences between sexes, and morphologically males and females have similar visual acuity (Fig. 5). In the absence of airborne cues, there is qualitative evidence that females are more attracted moving targets than stationary ones (Sippell and Brown, 1953; Kennedy, 1940; Wood and Wright, 1968) possibly because motion itself contributes to the perception of the target as a host. Females can only produce eggs after a blood meal (Beklemishev, 1940; Valzania et al., 2019), and have increased navigational gains toward tall dark environmental features representative of a host until the ingestion of a blood meal (Barredo et al., 2022). In contrast, males have no physiological need for a blood meal, but gather in the vicinity of a host where they intercept potential mates (Hartberg, 1971; Jones and Pilitt, 1973). Our finding that males stop approaching a moving target aligns with previous observations that motion reduces target attractiveness in males (Muir et al., 1992). This is ethologically intuitive; males risk an antagonistic interaction with little to gain when approaching a moving target.

**Figure 5.**
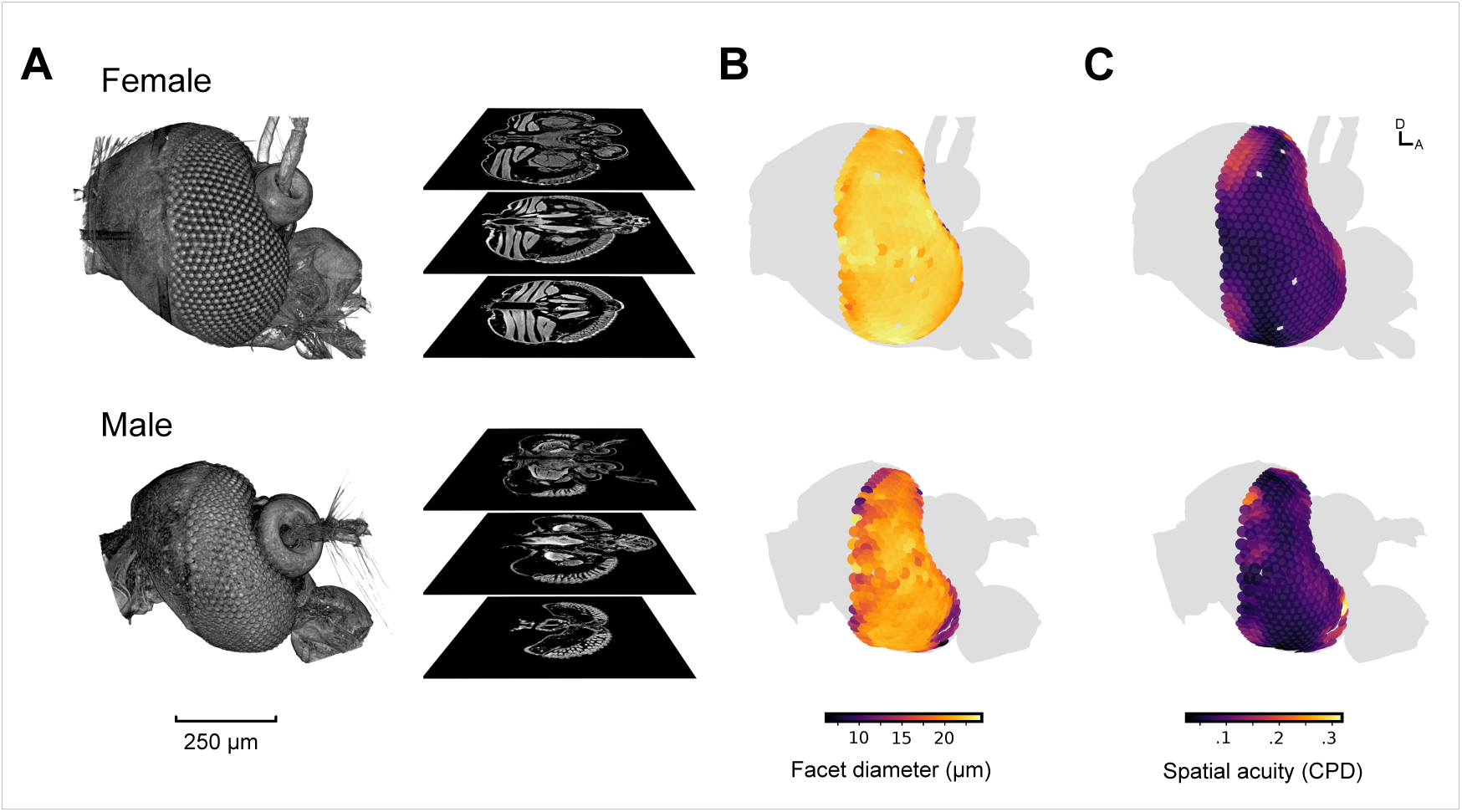
Male and female *Ae. aegypti* have similar visual acuity. **A:** µCT reconstruction of the head of a female (top row) and a male (bottom row) *Ae. aegypti*, showing observed regional variation in estimated facet diameter and spatial acuity. From left to right, three-dimensional rendering of the head followed by representative µCT cross-sections. **B** Heatmaps represent the distribution of skewness-adjusted facet diameters (left) and **C** the distribution of estimated spatial acuity across the compound eye. A: anterior, D: dorsal

In order to perceive a target’s motion, mosquitoes first need to disentangle it from their self-generated optic flow, a computationally demanding task with the sensory information available to a mosquito while in flight. During free flight, the motion of a static feature along the mosquito’s retina should, to some degree, be coherent with its motor output, but the optic flow patterns of a moving feature will be the combination of the mosquito’s motor output, drift in space, and motion of the target itself. Ommatidia, the functional units of insect eyes, detect these patterns as correlated changes in contrast across neighboring units on the retina. A feature’s retinal slip and consequential changes in luminance to motion-detecting units could be due to either a gust of wind blowing a mosquito off course, the target itself moving, or general drift in the mosquito’s flight motor. Based on the timing, axis, and polarity of the luminance change detected, a flying insect produces a corrective steering response aimed at stabilizing a feature onto a particular region of its retina, or to correct for unexpected panoramic motion (Borst, 2000; Hassenstein and Reichardt, 1958). How mosquitoes process optic flow information to gate approach or avoidance responses between the sexes remains to be investigated in detail.

A typical mechanism for positioning a target on the retina uses a simple control heuristic that may not compute the target’s motion at all. Typically, insects generate motor commands simply to center a target on a setpoint position on their retina (Fabian et al., 2018; Mischiati et al., 2015). In the absence of any other sensory information or motor feedback, a navigation strategy that only corrects a proportional error, in this case the offset of the feature from the desired setpoint on the retina, cannot resolve whether the feature has moved due to the mosquito’s own drift or due to the feature itself moving. Knowing that a tall feature is in motion requires a mosquito to compare information from multiple visual pathways. A moving feature having negative valence in one sex and a navigational strategy that centers it in another suggests mosquitoes possess the neural circuitry to determine external motion.

Information from the coupled expansion of the bar on the mosquito’s retina, combined with its change in position, could explain the fundamental differences between bar tracking between mosquitoes tethered with an unrestrained yaw axis girded in virtual reality and those reacting to projected stimuli in free flight. It is possible that mosquitoes require simultaneous comparison of their own motor state, feature expansion, and retinal slip in order to resolve whether a feature itself is in motion. This would explain why dark moving features in magnetically tethered virtual reality, a case with no visual expansion, could be interpreted as having a positive valence for male orientation and tracking, but in free-flight the target motion elicits an avoidance response. Alternatively, motion may elicit avoidance responses at a threshold projected width onto the male’s retina. In the virtual reality system, a male mosquito may not be able to resolve that the bar’s motion is not external to the mosquito’s own motion. From the mosquito’s point of view, it may be adjusting its steering responses to correct for drifting off a straight course. This incongruency between free-flight and partially-restrained preparations calls for a layer of caution for the interpretation of our understanding of organismal navigation generally.

While the motion of a visual target deters a male’s approach, we cannot yet determine whether motion adds an additional layer of attraction for females. Females approach a target regardless of whether or not it is in motion, but we are unable to resolve if a moving target is more attractive in this context. Previous work has suggested that female *Ae. aegypti* are more attracted to moving features than static ones when presented only with the visual of an anesthetized versus moving deer mouse (Sippell and Brown, 1953). Previous work also has suggested that females perform stronger turning responses toward moving targets compared to static ones (Kennedy, 1940). At close range, female *Ae. aegypti* evaluate a breadth of cues before landing on a potential host, including *CO*_2_ (Van Breugel et al., 2015), skin volatiles (Giraldo et al., 2023; De Obaldia et al., 2022), heat (Greppi et al., 2020; Giraldo et al., 2023), and skin wavelength content (Alonso San Alberto et al., 2022). Target motion may also strengthen the multi-modal signal that a female is approaching a host, rather than an inanimate feature.

Even though males may exhibit avoidance responses to object motion compared to females, the observation that the females’ flight trajectory is easily manipulated by moving targets may have important implications for visual lure design. The use of motion, either to mimic features of host motion or co-option of wide field motion steering responses, may be able to inform and improve field traps along with other modifications with better catch efficiency based on visual processing (Bidlingmayer, 1994). Because we demonstrate that mosquitoes can perceive motion, further investigation into whether host-like motion is more attractive to female mosquitoes than static features has the potential to influence control efforts. For example, co-opting optomotor responses, which normally correct for panoramic motion, may aid in luring mosquitoes into low-power, easily deployable field traps. Moreover, full characterization of mosquito visual processing, particularly the precise filtering properties that distinguish ego-motion from external target movement, holds promise for identifying molecular or genetic targets that compromise mosquito vision. By hijacking the flight control circuitry of the mosquito brain, we may be able to guide insects more reliably into trapping systems, ultimately improving vector control efforts with visual ecology-informed engineering strategies.

## AUTHOR CONTRIBUTIONS

Conceptualization: CR, SDS, JT, JAR Methodology: CR, SDS, RQ Software: CR, SDS, AW Validation: CR, SDS, SD Formal Analysis:CR, SDS, RQ Investigation:CR, SDS, RQ, AW, SD Resources: JAR, JT Data Curation: CR, SDS, RQ Writing-original draft: CR, SDS Visualization: CR, SDS, RQ Supervision: JAR, JT Project Administration: JAR Funding Acquisition: JAR, JT, SDS

### ACKNOWLEDGEMENTS

We would like to acknowledge Binh Nguyen for assistance with mosquito rearing and members of the Riffell Lab, especially Saumya Gupta, Cleopatra Pimienta, Takuro Ohashi, Adam Blake, and Genevieve Tauxe for comments and suggestions during the undertaking of this work. Omar Akbari and Fangying Chen additionally provided valuable comments.

## DATA AVAILABILITY

All data along with figure-generating code will be made publicly available via Dryad upon publication.

## FUNDING

This project was funded by the Air Force Office of Scientific Research under grants FA9550-20-1-0422 (JAR) and AWD-004055-G4 (JAR and JT); National Institutes of Health under grants R01AI175152 and R01AI148300 (JAR); the National Science Foundation, IOS-1750833 (JT); and the Air Force Research Laboratories grant FA8651-25-2-0004 (SDS).

## COMPETING INTERESTS

The authors have no competing or financial interests.

## Notes

### Competing Interest Statement

The authors have declared no competing interest.

